# Biodiversity clock and conservation triangle: Integrative platform for biodiversity monitoring, evaluation, and preemptive conservation intervention

**DOI:** 10.1101/2019.12.16.877969

**Authors:** R Jaishanker, M Vishnu, Sajeev C Rajan, N.P Sooraj, K Athira, V Sarojkumar, Lijimol Dominic, Subin John Mathew, Anjaly Unnikrishnan, Vinay Kumar Dadhwal

**Affiliations:** C V Raman Laboratory of Ecological Informatics, Indian Institute of Information Technology and Management-Kerala, Thiruvananthapuram, Kerala, India-695581; Indian Institute of Space Science and Technology, Trivandrum, Kerala, India

**Keywords:** Biodiversity, IUCN conservation status, CBD, Post 2020 biodiversity framework

## Abstract

Business-as-usual is no more an option on the table for biodiversity conservation. Disruptive transformation at both policy and polity levels are pressing needs. The possibilities presented by the current wave of information and communication technology can act as travelators to meet the conservation targets. Here, we introduce twin concepts of biodiversity clock and conservation triangle that posit as convergence plane to seamlessly consolidate ongoing discrete efforts and convey real-time biodiversity information in a lucid schematic form. In its present form, the biodiversity clock depicts 12 ecological and 6 biophysical components. The universal consistency in clock-reading facilitates the biodiversity clock to be read and interpreted identically across the world. A ternary plot of the International Union of Conservation of Nature (IUCN) species conservation status is presented as the conservation triangle. Together, the biodiversity clock and the conservation triangle are invaluable in strategizing biodiversity conservation, post-2020. Leveraged smartly, they make possible pre-emptive intervention for biodiversity conservation.

## Introduction

Biological diversity is a strategic investment made by nature that ensures perpetual competition and progressive improvement of life forms. Individual species, like entrepreneurs, scout for opportunities (niches) within and beyond their native habitats. Successful entrepreneurs acquire more niches, gain a competitive edge, and become keystone assets. The synchronized orchestra of different life forms generates value-added (ecosystem) services. As an immaculate bandmaster, nature has designed seamless ways to ensure *cooperative-competition* among different species. However, unlike the notion of collective action in human societies [1], individual cost-benefits in the natural schema are conjoined with that of the whole. The splice ensures no free lunch and preempts a proprietary claim to the ecosystem services by any species. All species have to contribute in order to enjoy the benefits of value-added services. The system satiates both the micro and macro contributors. While the former gets a habitat, the status of the latter (keystones) is imaginary, sans the former.

The total productivity of an ecosystem is linked to the individual productivity of all constituent species and is governed by the limiting factor resource [2]. Nature has designed the system in such a way as to predominantly expend the value-added services, *in-situ*. The design of activities (biotic-biotic, biotic-abiotic and abiotic-abiotic interactions) and the spatio-temporal distribution of resources within (eco) systems, help to generate unique services. Within a stable structural ensemble, ecosystems remain sustainable and productive. Here too, the ingenuity of the bandmaster is on display. Albeit stable, structural ensembles are not rigid. They are amenable to gradual changes, which in-turn coerce the vying species to be on the perpetual look-out for emerging niches.

One among the 8.7 million entrepreneurs outgrew and has laid siege to all the other species. The authors and the reader are members of this species, which is predisposed to hoard resources, much in excess of what is needed and amass them from far beyond their surroundings. *Ex-situ* consumption of ecosystem services and mindless activities that led to accelerated changes of the structural ensemble; summarizes the actions of this species. The hastened rate of structural changes disproportionately put other species, at a disadvantage.

While we are yet to arrive at a way to express the hardships of other species, increased frequency of droughts, deluges, and emerging epidemics are the obvious harbingers of ecosystem disruption, that stems from our action. They sting and undermine human existence, too [3–5]. Hence, far more than other species, we should be concerned about their survival. Stated otherwise, human endeavor to conserve biological diversity has a latent element of self-centeredness. However, the conjoined benefit-sharing paradigm forbids man to take pride in the self-interest as his magnanimity. We are bound to the rules of *competitive-cooperation* and can fall back from our responsibility only at our own risk.

*Sapiens* is a responsible species. Much before it became a *felt need*, the importance of environment and wildlife conservation echoed in the corridors of power. President Theodore Roosevelt was famous for his executive orders in natural resources conservation and wildlife protection [6]. Nonetheless, it took more than six decades since the conservation concerns of President Roosevelt [7, 8] for the world to rise collectively for the protection of the environment and conservation of species. The United Nations conference on the human environment in Stockholm in 1972 was a landmark event that triggered a series of collective global efforts on environmental issues [9, 10]. The constitution of the International Union of Conservation of Nature (IUCN) in 1958 and the publication of ‘Silent Spring’ [11] are two towering milestones that, along with the other similar initiatives, helped to coalesce the discreet efforts, in Stockholm. Strangely, despite rapidly increasing awareness of the consequences, persistent *stultus* (foolish) behavior of the *sapiens* (wise), defies logic. With mechanized tools, no corner of the planet was beyond human reach. We pulled down the structural ensembles built over eons, in a couple of centuries. The dilapidated structures (habitats) [12–14], became permeable to alien species that outsmarted and decreased the native diversity [15]. We have shoved more than a third of all known species into unsafe zones where they face an enhanced threat of extinction [16].

Meanwhile, the international chorus for biodiversity conservation and sustainable development progressively gained decibel at Nairobi, Rio de Janeiro, Johannesburg, and Paris [17]. The success of the Montreal Protocol [18] lends hope for concerted action in addressing the environmental challenges. The Convention on Biological Diversity (CBD) identified the conservation of biological diversity as a global human concern and presented a legally binding framework. Within the CBD, the strategic plan for biodiversity encompasses 20 time-bound targets - the Aichi Biodiversity Targets [19], which concurrently feed into the Sustainable Development Goals (SDGs) [20].

Protocols of practical estimation of biodiversity constitute the basis of the assessment of progress to the Aichi targets. Documenting diversity and representing its temporal dynamics is fundamental to the Aichi targets [21, 22]. Although the option to determine the approach to assess progress towards implementation of the strategic plan for biodiversity (from among quantitative indicators, expert opinion, stakeholder consultation, and case studies), rests with individual *Parties*; the advisory to ensure repeatability is a veiled emphasis in favor of quantitative indicators [19]. While a well-conceived indicator also doubles as good surrogate to evaluate the success of conservation interventions [23], the inconsistent use of indicators [24] and the heterogeneous data compilation at micro-levels [25] by the *Parties*, impede constructive spatial aggregation and temporal analysis. Hence, in the penultimate year of the international decade of biodiversity, we are still struggling with Tier II and III indicators for multiple Aichi targets [20]. All is not lost. The present wave of Information and Communication Technology (ICT) gifts us with travelators to meet Aichi commitments. A plethora of online ecological and ecophysical database was washed ashore during the first four waves of ICT. Searchable IUCN Red data book (www. https://www.iucnredlist.org/), Global Invasive Species Database (http://www.iucngisd.org/gisd/), Plant Trait database (https://www.try-db.org/TryWeb/Home.php) are few representatives of the former. Group on Earth Observation Biodiversity Observation Network (GEO BON) (https://geobon.org/), PROBA-V (https://proba-v-mep.esa.int/applications/time-series-viewer/app/app.html), Earth Systems Research Laboratory, National Oceanic and Atmospheric Administration (ESRL, NOAA) (https://www.esrl.noaa.gov/psd/data/gridded), are instances of the ecophysical dataset. Although the challenge of spatial scalability has been addressed to a fair extent in the ecophysical data set, the ecological (and biodiversity) data are still plagued by heterogeneity and micro-scales. The present disruptive wave offers an unparalleled capability to collect, handle, and analyze ecophysical data for real-time representation and preemptive interventions for biodiversity conservation.

Here, we introduce a biodiversity clock and conservation triangle that can consolidate multiple dimensions to represent the biological diversity in a landscape. The duo of the clock and triangle acts as a foundation upon which an intelligent system can be built to autonomously assimilate and depict biological and ecophysical data, and keep a tab on its temporal changes.

## Materials and Methods

For the sake of clarity, we present the biodiversity clock and conservation triangle with a case study in two sacred groves in Kerala, India. Sacred groves are vibrant vestiges of erstwhile forest landscapes [26, 27], and are ideal for biodiversity studies. They are reported in many countries [28]. In India, 40,000 hectors (ha) of land are revered as sacred groves. More than 2000 sacred groves pepper the landscape of Kerala and occupy a geographic area of more than 500 ha [29, 30].

### Study Sites

Two sacred groves, *Kavil Shree Maheswar Ashramam* (KSA) and *Edayilekkadu Kavu* (EDK), in Nedumangad (Trivandrum) and Thrikkaripur (Kasargod) in Kerala, India were selected for the present study. While the former is situated in the foothills of the Western Ghats and spread over 36000 m^2^, the latter is a coastal grove that spans 40000 m^2^ (Fig 1). Situated in the southwestern side of the Indian peninsula, Kerala enjoys a tropical climate. Both the groves harbor predominantly evergreen tree species.

**Fig 1.**
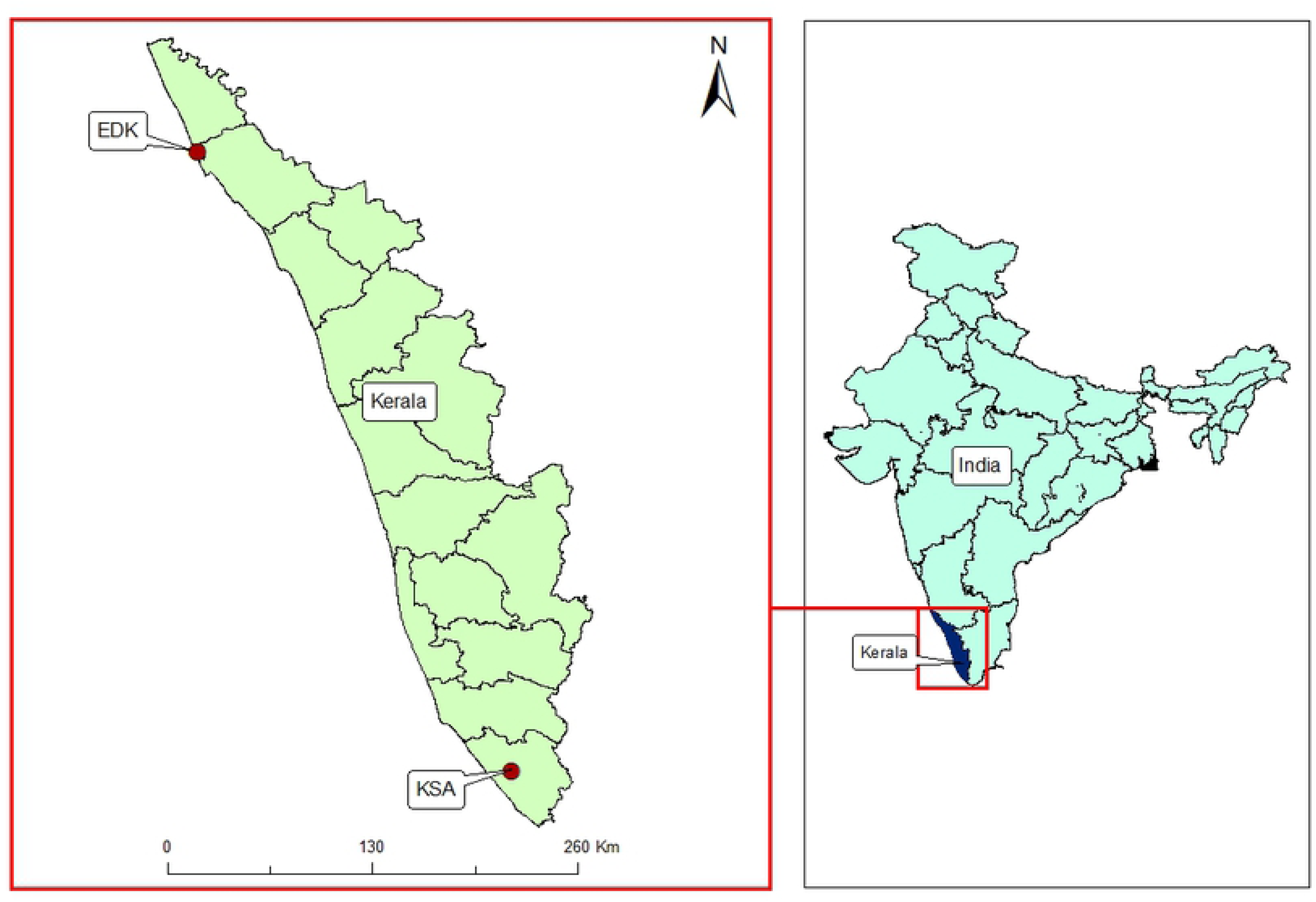
Map showing two sacred groves KSA and EDK.

### Field Survey

Field surveys were carried out in February 2019. We confined our study to tree species and included only those that measured a minimum 1-foot diameter at breast height (DBH). Since the extent of the sacred groves was small, total enumeration was adopted for the tree inventory. Irrespective of *habit,* the number of invasive plant species within the groves were recorded. The approximate length and width of each grove from its center were recorded as the semi-major and semi-minor axis (a, b) in Kilometer (km), respectively. The coordinates of the geometric center of each grove were also noted.

The spreadsheet template for ground data collection used for this study is shared as S1 File. The technical simplicity of the attributes in the spreadsheet was purposely ensured for easy replication. The canopy gap at each site was estimated from hemispherical images (Fig 2) taken using a fisheye lens mounted on a smart-phone, held perpendicular to and 1 meter (m) above the ground. Hemispherical images were taken at randomly selected locations within each grove to ensure representative coverage of the canopy gap (CNG). They were sequentially analyzed using R package - *Sky* [31] to determine the mean CNG (%) of the respective sites.

**Fig 2.**
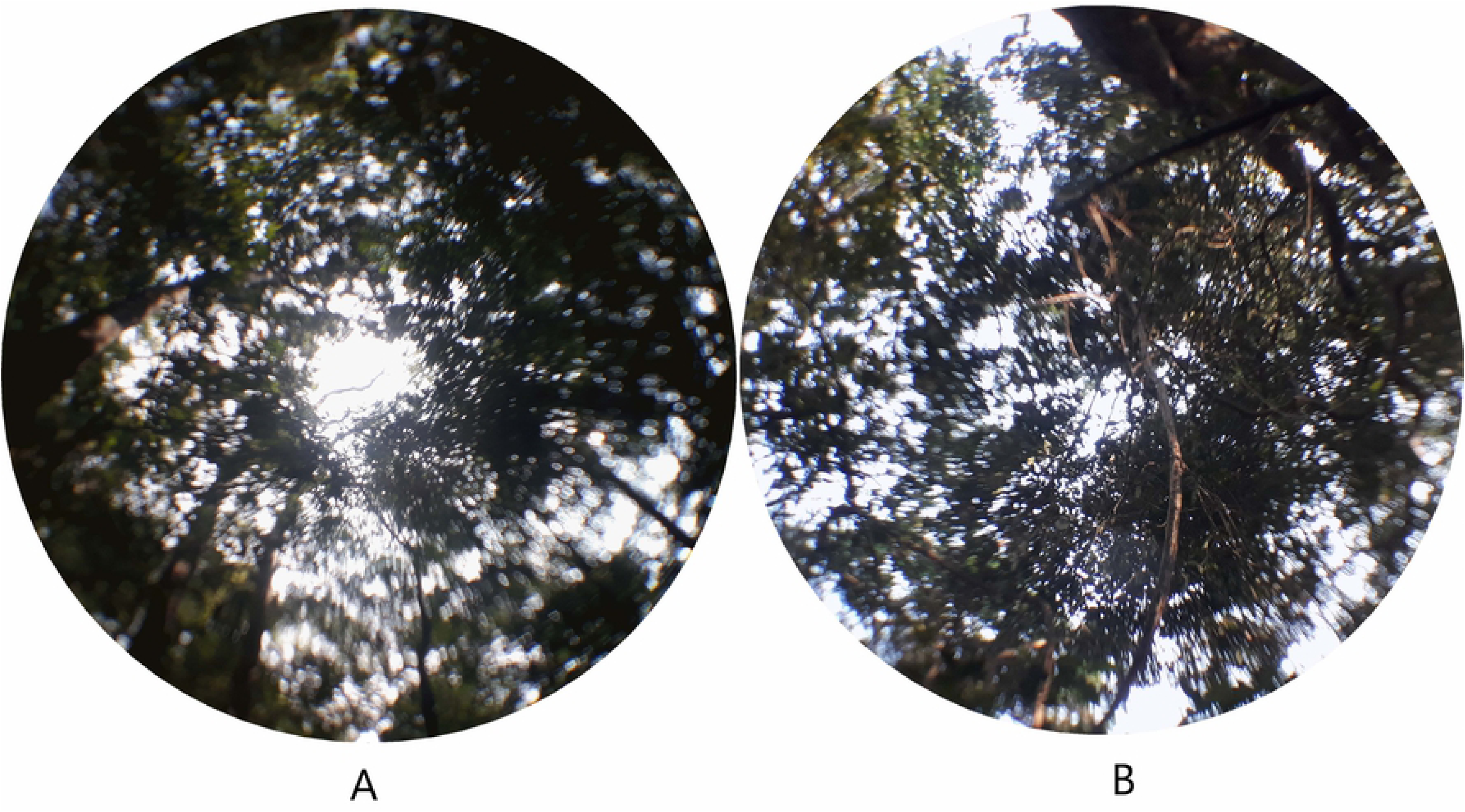
Representative hemispherical images of (A) KSA and (B) EDK.

### Data Processing

A system to generate the biodiversity clock and conservation triangle was developed in java platform. The populated data template for each site was uploaded to the system to generate its respective biodiversity clock and conservation triangle. The ensuing sub-sections describe the backend data processing involved.

#### Species Classification based on IUCN status

The system automatically classifies individual tree species in the uploaded dataset based on their IUCN conservation status [32] into the following six categories: Least Concern (LC), Near Threatened (NT), Vulnerable (VU), Endangered (EN), Critically Endangered (CR), Extinct in the wild (EW), and Extinct (EX). For those species, which are either ‘not evaluated’ or reported ‘data deficient’ by the IUCN, the system prompts the user to either accept the default class, Least Concern (LC) or to choose from the other IUCN classes. However, the submission of a valid reference is mandatory to exercise the latter option. The six groups (for practical purposes, 5) were subsequently regrouped into three categories, *viz*. LC, VU, and EN. While the LC and VU classes from the first iteration remain unaltered, CR, EW, and EX were merged with the EN class.

#### Biodiversity Description

Subsequent to the successful classification of tree species, the system estimates the species richness, Shannon effective number (Macarthur 1965), the proportion of LC *(PLC)*, VU *(PVU)* and EN *(PEN)* species, the proportion of endemic, and keystone species in each of the three IUCN conservation classes, respectively. For want of a global database, the status of endemic, and keystone species are currently set as user entered attributes. The system searches the public domain of the Invasive Species Specialist Group (ISSG) (www. iisg.org) to ascertain the invasive status of each species in the data template.

### Biodiversity Clock

#### Biodiversity and Ecophysical Components

Twelve biological and six eco-physical variables of each sacred groves were represented as clock graphs. Species richness *(S)*, Shannon effective number *(Es)*, proportion of invasive species *(PIN)*, proportion of LC *(PLC)*, proportion of VU *(PVU)*, proportion of EN *(PEN)* species, proportion of endemic species in LC *(PeLC)*, proportion of keystone species in LC *(PkLC)*, proportion of endemic species in VU *(PeVU)*, proportion of keystone species in VU *(PkVU)*, proportion of endemic species in EN *(PeEN)*, proportion of keystone species in EN *(PkEN)*, were the biodiversity variables depicted.

The time of field measurement, indicated as month:year (mm:yyyy), and geographic coordinates *(Lat:Long)* of the center of the study site are directly read from the database. The extent of the study area was expressed as the semi-major and semi-minor axis *(a,b)* expressed in kilometer (km). The maximum and minimum elevation within the study area was estimated as the mean elevation value of 20% randomly selected pixels within a bounding box of dimensions *a,b,* and read from the Google Earth. These constitute the eco-physical attributes displayed in the biodiversity clock. In addition, the system is designed to store the coordinates of each uploaded data template into a database that builds up over time to facilitate regional aggregation.

#### Clock Reading

The key to interpreting the biodiversity clock is illustrated in Fig 3. Biodiversity and biophysical variables are denoted in blue and red fonts, respectively. The description of each component with its representation is given as S2 File.

**Fig 3.**
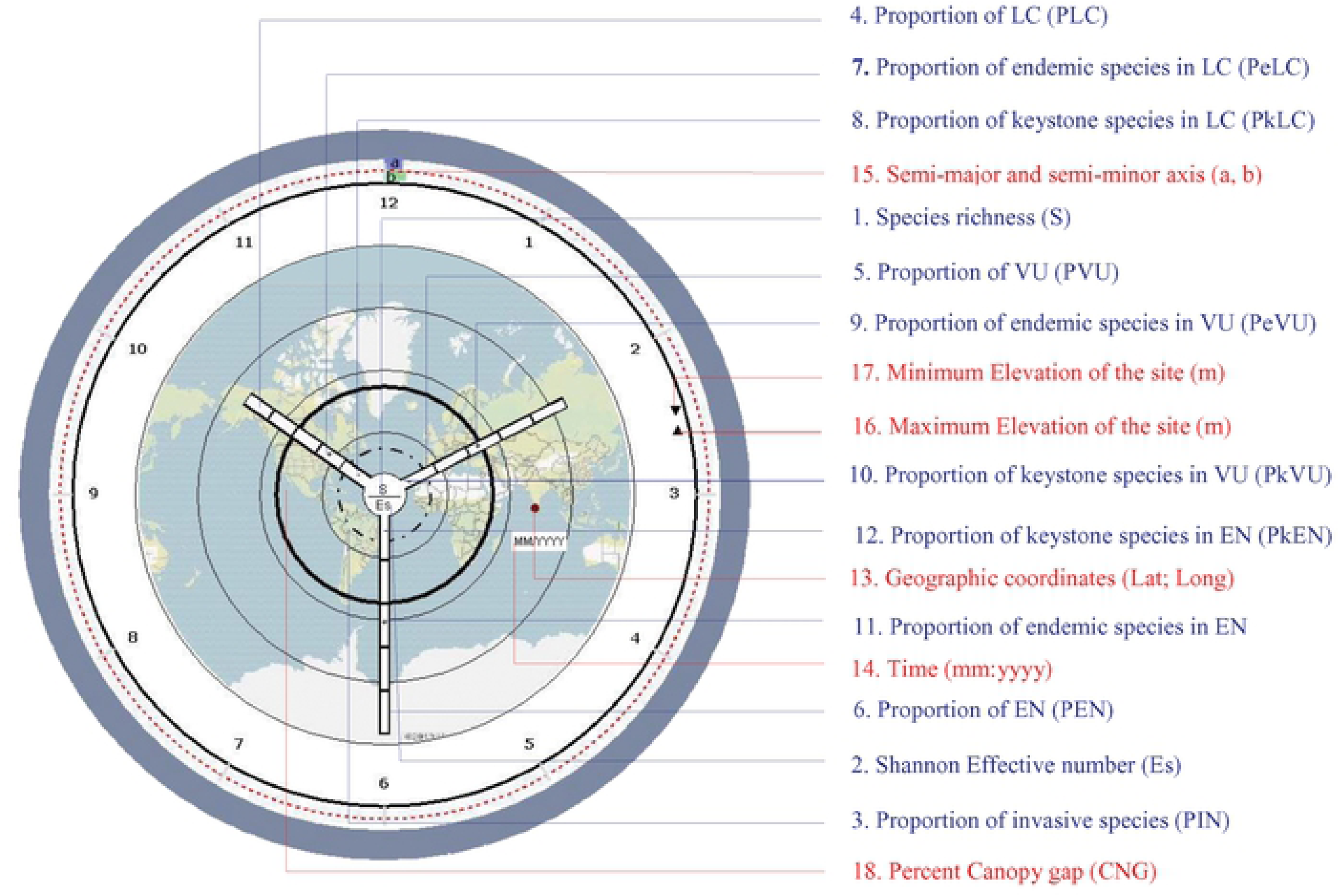
Interpretation key for biodiversity clock. Description and scheme of representation are provided in S2 File.

#### Conservation Status

The three classes, LC, VU, and EN, were depicted on the hour, minute, and second hand of the biodiversity clock, respectively. The time displayed on the biodiversity clock was arrived at by rescaling the proportion of LC, VU, and EN classes to 0 to 360 degrees. The angle subtended by each hand is the product of its corresponding proportion with 2π, measured clockwise from a hypothetical radius to the 12-hour mark on the dial.

The conversion of population proportion to time representation is explained below.

Let S be the total number of tree species, PLC, PVU, and PEN be the corresponding proportions of the categories *viz*. LC, VU, and EN such that PLC + PVU + PEN = 1

If PLC, PVU, and PEN are represented by the hour, minute, and second hand of a clock respectively; then

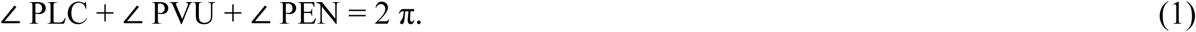

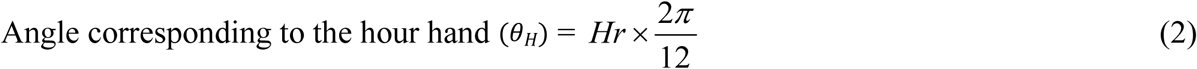

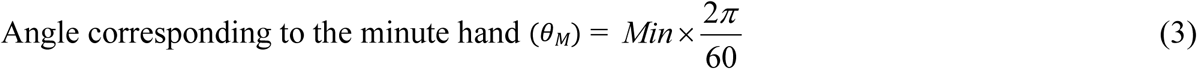

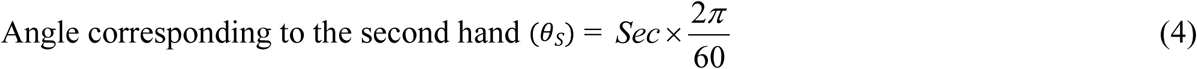

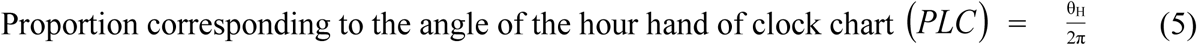

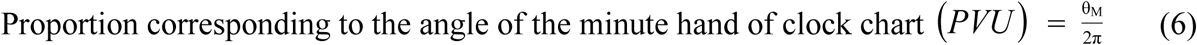

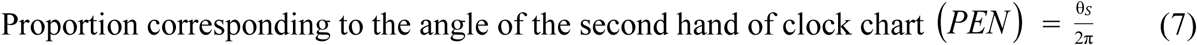

#### Biodiversity Variables

The Species richness *(S)* and the Shannon effective number *(Es)* are displayed as the numerator and denominator of the fraction indicated on the head of the central pivot of the clock. The proportion of endemic and keystone species in each of the three IUCN conservation classes are depicted in the corresponding hand of the clock. For ease of interpretation, each hand is divided into 5 equal segments, and the proportions of endemic and keystone species are indicated with plus (+) and minus (-) symbol in corresponding segments of the respective hand. The proportion of endemic *(PeLC)* and keystone *(PkLC)* species in the LC class is indicated with + and − symbols in the appropriate segment of the hour hand. The proportion of invasive species in the population *(PIN)* is denoted as a dashed circle positioned appropriately on the dial, which is indicatively divided into five equiproportional concentric circles. The pinhead is scaled to 5%, and lower proportions are denoted as dashed lines on it.

#### Eco-physical Variables

The time of field measurement is displayed in the time window, and the geographic location of the study area is indicated with a dot symbol on a physical map, set as the background of the dial. The bezel of the clock is split into two concentric rings. The outer and inner rings denote the semi-major and semi-minor axis *(a, b),* respectively. Both rings are divided into 12 sections. Each segment is scaled to represent a ground distance in km with base 2 and increases geometrically as 2, 2^2^, 2^3^, 2^4^, …. 2^12^ km, respectively. The first segment of both *a* and *b* denotes 2 km, the second; 4 km and the twelveth; 4096 Km. The spatial extent of the study site is represented by highlighting the corresponding *a* and *b* segments on the bezel.

The maximum and minimum elevation of the study sites are depicted with a triangle (▴) and inverted triangle (▾) symbol, respectively, along the rim of the clock. A modified Fibonacci series; 0, 50, 100, 150, 250, 400, 650, 1050, 1700, 2750, 4450, 7200, 11650 was used as the reference scale. Read from the second element (50), the series corresponds to 1,2,3,4,5,6 ….., 12- hour marks on the dial, respectively. Sea level and the highest possible elevation above the Mean Sea Level MSL (11650 m) fall on the 12-hour mark. Since the highest land elevation above MSL on the Earth is 8848 m, chances of any ambiguity in representation are nil. The same schema is followed to depict the surface below sea level. However, the maximum and minimum depth of surface below MSL is depicted with Delta (Δ) and Nabla symbols (∇), respectively. The maximum possible depth that can be depicted on the clock graph is −11650 m.

The five equiproportional concentric circles on the dial act as the reference frame to interpret the *CNG* of the study site. The % *CNG* of the study site is highlighted as an appropriately positioned solid circle on the dial.

#### Sensitivity Analysis

The angular sensitivity of the biodiversity clock to changes in the number of LC,VU, and EN classes was assessed against a reference basal population of tree-species (S). The analysis was carried out in two dimensions *viz.* a reduction of 1 species in the IUCN categories, and a shift of 1 species across the IUCN categories.

Let S denote the total number of tree species and θ_H_, θ_M_ and θ_S_ be the angle that corresponds to the hour, minute, and second hand of the biodiversity clock, respectively.

***Scenario 1***. Loss of 1species in the IUCN categories

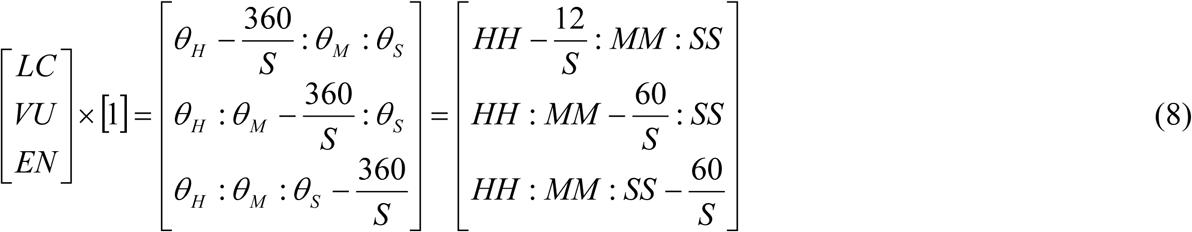

***Scenario 2***. A shift of 1 species across the IUCN categories.

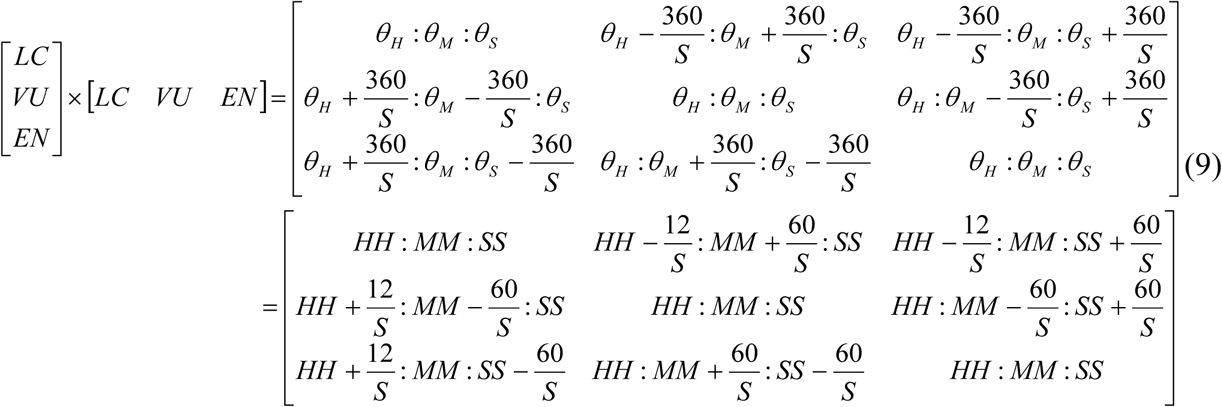

### Conservation Triangle

The proportion of tree species in the three IUCN conservation classes (*PLC, PVU, PEN*) were represented using a ternary plot to arrive at the conservation triangle. The number of tree species under each IUCN conservation class was normalized to 0 to 100. The generalized equation that describes the composition of species (X_R_) is

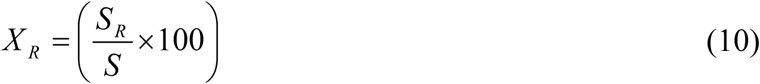

where *R* denotes the categories (LC, VU, and EN). *S_R_* is the number of species in respective categories and *S*is the total number of tree species.

*X_LC_ X_VU_* and *X_EN_* constitutes the three axes of the ternary plot. The corners of the triangle represent the saturated space of the corresponding classes. The apex (A) of the conservation triangle denotes 100% EN, 0% VU & 0% LC. The lower right (B) and left (C) vertices represents population with 100 % LC, 0% VU & 0% EN and 100% VU, 0% EN & 0% LC, respectively. The proportion of LC species increases from 0 to 100 along 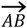. Similarly, the proportion of VU and EN tree species increases from 0 to100 from 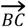 and 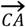, respectively. The R code used to create the conservation triangle is provided in S3 File.

## Results

### Biological and Ecophysical variables

The biodiversity and eco-physical variables of the sacred groves, computed by the system, were presented in Table 1. The species richness of KSA (48) was more than twice that of EDK (22). KSA and EDK had an equivalent tree diversity as that of communities with 26 and 13 equally common tree species, respectively. The proportion of LC tree species in KSA and EDK was 0.8 and 0.54, respectively. While 0.12 and 0.32 tree species in KSA and EDK belonged to VU class, respectively. The proportion of EN class tree species in EDK (0.14) was higher than that in KSA (0.08). *Mikania micrantha* was the only invasive plant species observed along the fringes of KSA. No invasive plants were observed in EDK.

**Table 1.** Biodiversity and Eco-Physical Variables of the study sites.

| SNo | Variable | KSA | EDK |
| --- | --- | --- | --- |
| **Biodiversity** |  |  |  |
| 1 | Species richness (*S*) | 48 | 22 |
| 2 | Shannon Effective number (*Es*) | 26 | 13 |
| 3 | Proportion of invasive species (*PIN*) | 0.02 | 0 |
| 4 | Proportion of LC (*PLC*) | 0.80 | 0.54 |
| 5 | Proportion of VU (*PVU*) | 0.12 | 0.32 |
| 6 | Proportion of EN (*PEN*) | 0.08 | 0.14 |
| 7 | Proportion of endemic species in LC (PeLC) | 0.08 | 0.25 |
| 8 | Proportion of keystone species in LC (PkLC) | 0.03 | 0.08 |
| 9 | proportion of endemic species in VU (PeVU) | 0.5 | 0.29 |
| 10 | Proportion of keystone species in VU (PkVU) | 0 | 0 |
| 11 | Proportion of endemic species in EN (PeEN) | 0.5 | 0.67 |
| 12 | Proportion of keystone species in EN (PkEN) | 0 | 0 |
| **Eco-physical** |  |  |  |
| 13 | Geographic coordinates (*Lat; Long*) | 8.60; 77.95 | 12.12; 75.16 |
| 14 | Time (*MM: YYYY*) | 02/2019 | 02/2019 |
| 15 | Semi-major and semi-minor axis (a, b) | 0.12, 0.075 | 0.135, 0.085 |
| 16 | Maximum Elevation of the site (m) (▲) | 132 | 13 |
| 17 | Minimum Elevation of the site (m) (▼) | 123 | 7 |
| 18 | % Canopy openness (*CNG*) | 22 | 14 |

The presence of endemic species within the groves was expressed as its proportion in the respective IUCN classes (LC, VU, and EN). The proportion of endemic tree species in EN class (*PeEN)* at KSA and EDK was 0.5 and 0.67, respectively. The proportion of endemic tree species in the VU class (PeVU) at KSA and EDK was 0.5 and 0.29, respectively. One and two-eighths LC tree species in KSA and EDK, respectively, were endemic. *Ficus religiosa* and *Ficus benghalensis* were the keystone tree species observed at KSA and EDK, respectively.

The angle subtended by the upright and inverted triangles with the pinhead and their respective positions on the clock dial throws light on the surface relief of the study sites. With 14% CNG, EDK had a more closed canopy than KSA (22%). With the highest elevation, 13 m EDK is almost at the sea level, whereas KSA is situated about 132 m above MSL (Table 1). The list of tree species, in the two sacred groves along with their conservation status, are presented in S4 File.

### Biodiversity clock

The tree diversity in the two sacred groves, KSA, and EDK, was figuratively represented as biodiversity clocks in Figs 4 and 5, respectively. The time on the clock corresponds to the proportion of the tree species in the IUCN conservation classes (LC, VU, and EN), represented by the hour, minute, and second hand, respectively. The hour hand between 9 and 10-hour mark on the dial corresponds to an angle of 280 degrees (equation 2 & 5) denotes 80% Least Concern (LC) tree species in KSA (Table 1). 12% Vulnerable (VU) tree species in KSA (Table 1) corresponds to an angle of 14 degrees subtended by the minute hand (equation 3 & 6). 8% Endangered (EN) tree species in KSA (Table 1) corresponds to an angle of 09 degrees subtended by the second hand (equation 4 & 7). Hence the biodiversity clock of KSA in February 2019 reads 09 Hours, 08 minutes, and 05 seconds (Fig 4). With 54%, 32%, and 14% of LC, VU, and EN classes respectively, the biodiversity clock for EDK denotes 07 hours, 20 minutes, and 08 seconds (Fig 5).

**Fig 4.**
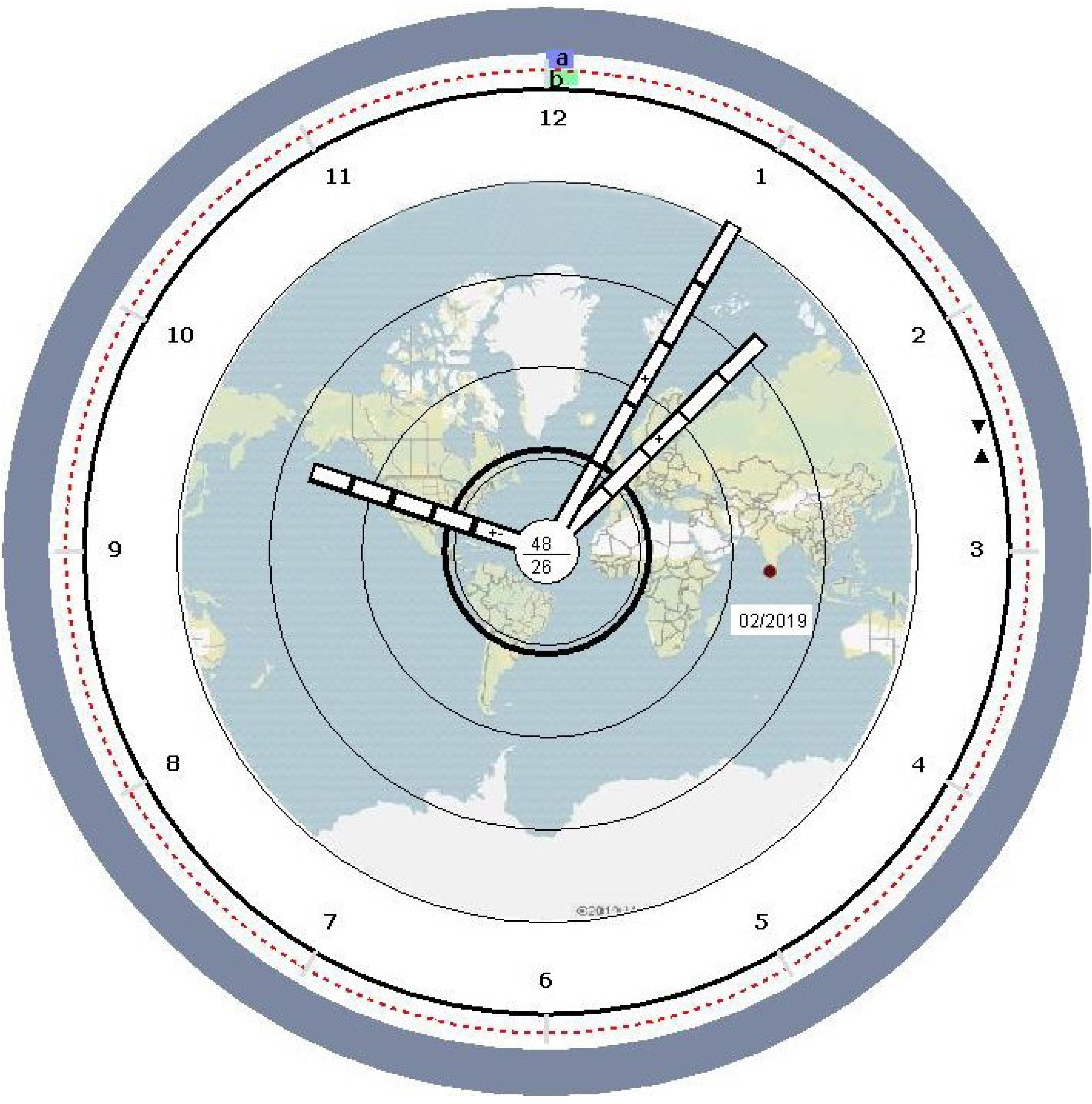
Biodiversity clock of study site - *Kavil Shree Maheswar Ashramam* (KSA), Kerala, India.

**Fig 5.**
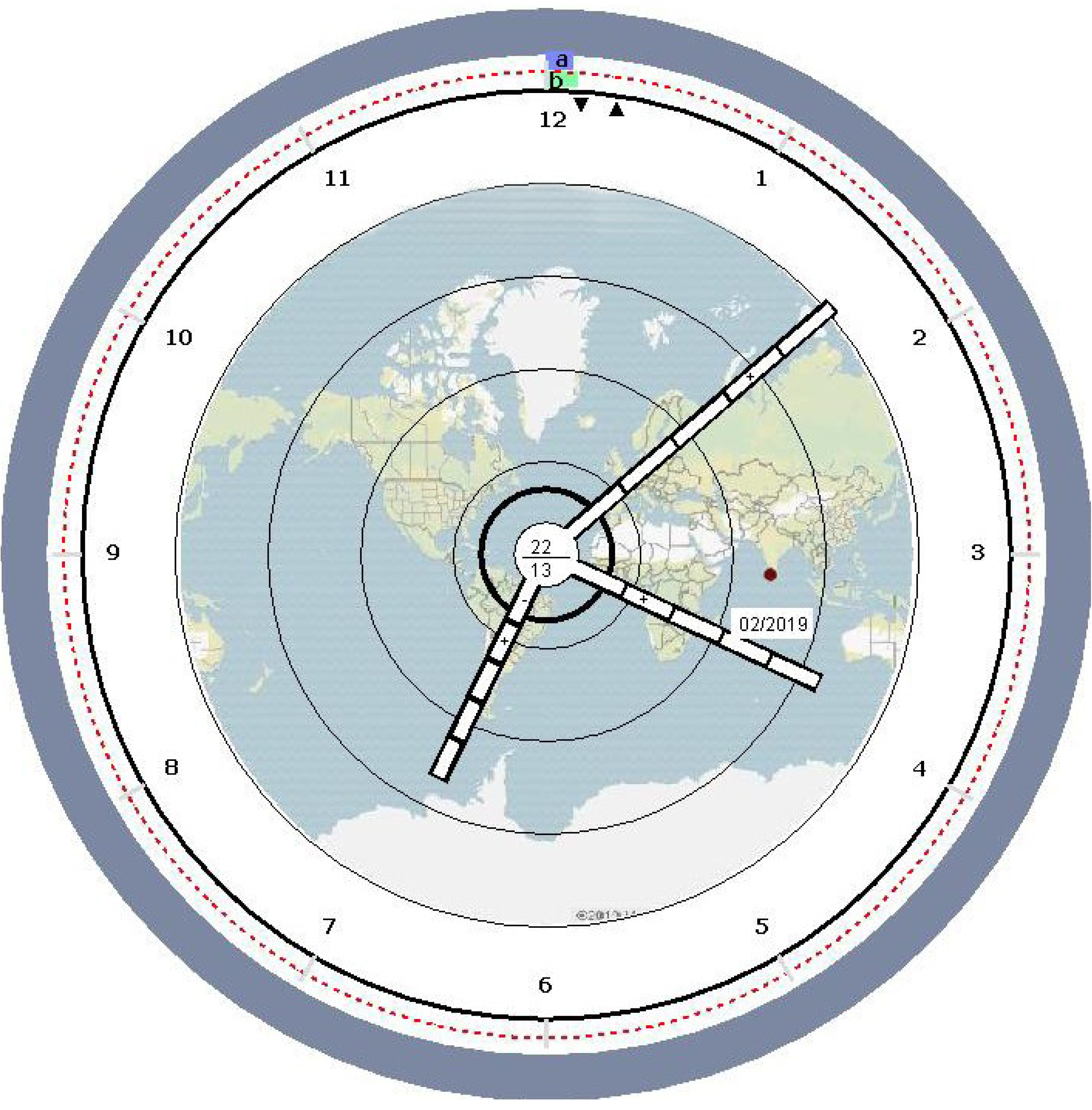
Biodiversity clock of study site - *Edayilekkadu Kavu* (EDK), Kerala, India.

Species richness (48 & 22; KSA & EDK) and Shannon effective numbers of the groves (26 & 13; KSA & EDK) are displayed as the numerator and denominator on the pinhead (Figs 4 and 5), respectively. 8%, 50%, and 50% tree species in the LC, VU, and EN classes in KSA respectively were endemic. 25%, 29%, and 67 % tree species in the LC, VU, and EN classes in EDK respectively were endemic. These were indicated with the + symbol in the appropriate segments of the hour, minute, and second hand, respectively.

The dashed line within the pinhead of the biodiversity clock of KSA indicates the *PIN*. The highlighted solid ring within the first and second concentric circles on the dial indicates 22 and 14 % *CNG* at KSA and EDK, respectively.

For each unit change in the number of species in LC, VU, and EN, the clock hands shift by 2π/S degree. Similarly, clock hands shift by 12/S degree for Hour hand and 60/S degree for Minutes and Second hands for each unit shift of species across LC, VU, and EN. The apportioning of the three clock hands to the respective IUCN species conservation category was based on the respective sensitivity. The hour-hand, representing LC, was less sensitive, as compared with the other two hands, which are identically sensitive.

### Conservation Triangle

The ternary plot of the proportion of tree species in three conservation classes LC,VU, and EN, respectively, was put forth as the conservation triangle (Fig 6). A ternary plot depicts three variables, which sum to a constant. The conservation triangle graphically depicts the ratio of the PLC:PVU:PEN as a unique position. In the present case, the constant was the total of PLC + PVU + PEN, within KSA and EDK, respectively. The conservation locus of each site KSA and EDK is the respective point within the plot, where the corresponding values of the proportion of LC,VU, and EN, intersects. The conservation locus of KSA with a predominantly high proportion of LC tree species (80%) falls in the green-hued zone of the conservation triangle. However, conservation locus of EDK with 32% VU and 14% EN tree species, is pegged in the transition zone between LC and VU classes.

**Fig 6.**
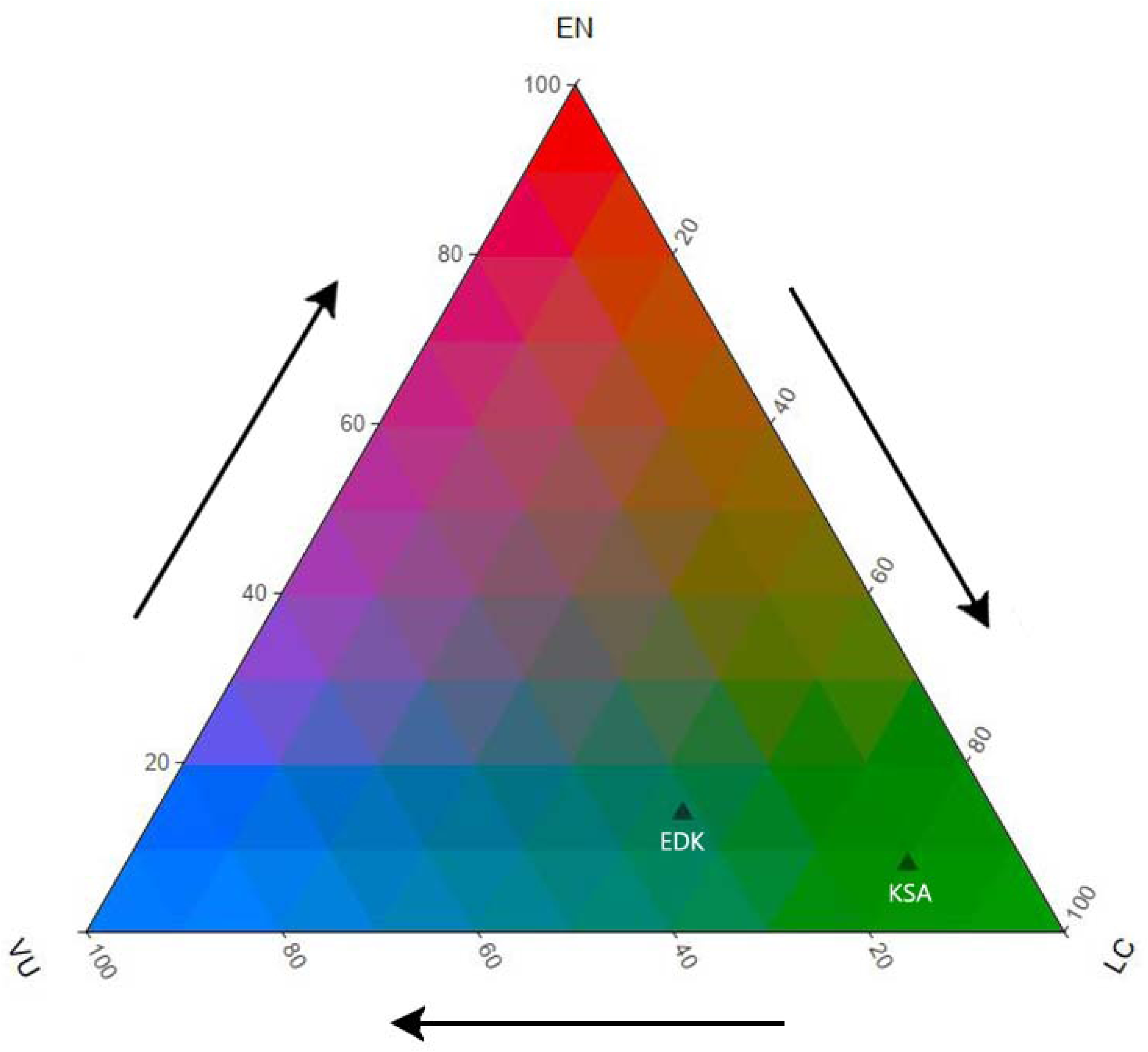
Conservation Triangle representing conservation locus of KSA and EDK.

## Discussion

### Biodiversity Clock

The clock - one of the oldest human inventions, stands out as perhaps the most uniformly interpreted device. The unit of time measurement and its schematic representation has remained above the cultural nuances that are otherwise evident in almost all other base units. Albeit clock reading is an acquired skill, dial with 3 hands is depicted and comprehended identically across all human cultures. The standard clock face has a short hour-hand, an intermediate minute-hand, and a long seconds-hand. It is customary to read a clock by first reading the hour, followed by the minute and second hands, respectively.

The inventor of the clock serendipitously stumbled on one of the core capabilities of the human brain - pattern recognition [34, 35], which endeared the template. The universal acceptability of the clock stems from its ability to organize and represent time information in a meaningful schema. The HH-MM-SS schema of time is a relational structure that enhances information comprehension. The *script/ frame* generated by a recurring schema of time [36] provides an unmatched option to convey semantics. We leveraged the clock schema to design the biodiversity clock.

The biodiversity clock is exemplified as case studies in two sacred groves, KSA and EDK, in Kerala, India. The clocks depicted in Figs 4 and 5 embody 12 biological and 6 physical dimensions respectively and provide a holistic representation of the two sacred groves. Sacred groves are socio-cultural edifices that help species conservation [26,28,37], and are ideal sites for long-term biodiversity studies. They are isolated patches of remnant vegetation [38], and occasionally entrapped in a rapidly urbanizing environment. Entrapped groves offer unparalleled prospects to study the effect of anthropogenic pressures on biodiversity. While KSA is predominantly in a rural environment and bound by plantations on all sides, EDK is an island of wilderness in a semi-urban setting.

The tree diversity at KSA and EDK were represented as respective biodiversity clocks. The time indicated by the hour hand of the clocks conveys the proportion of tree species within the LC category at the respective grove. The straight angle formed along the clockwise direction by either the hour, minute, or second hand, to an imaginary radius to the 12-hour mark, corresponds to a 50% proportion of the respective conservation class. A dial that reads 6 O’ clock or above comprises ≥ 50 % LC species. Biodiversity clocks for KSA and EDK showed 9 and 7 hours, respectively. It conveys the predominance of least concern tree species (*PLC*) at both sites. While *Calophyllum inophyllum, Phyllanthus emblica, Flacourtia montana* were the predominant LC tree species at KSA, *Holigarna arnottiana*, *Cinnamomum malabatrum*, and *Pongamia pinnata* were the most predominant LC tree species at EDK.

The right angle formed by either the hour, minute, or second hand with the imaginary radius to the 12-hour mark corresponds to one-fourth proportion of the respective conservation classes. The acute angles of the second hands of the respective biodiversity clock of KSA and EDK point to the lower proportion of endangered tree species (*PEN*) therein. While the minute hand of the biodiversity clock for KSA subtend an acute angle, that of EDK was obtuse. This conveys the relatively higher proportion of vulnerable tree species at EDK. The biodiversity clock for a population with equal proportions of the three IUCN conservation classes will show the time; 4 Hrs 40 Min and 0 Seconds. Since the biodiversity clock is bound by equation (1), an appropriate representation for the skewed proportion of LC, VU, and EN species is ensured. For instance, the biodiversity clock will not display the corresponding hand if either *PLC*, *PVU*, or *PEN* of a site equals zero. The biodiversity clock for a site with 100 % *PLC* will display only the hour angle, which will point to the 12-hour mark. Here the ∠ PLC is read 2 π and not zero. The additional variables represented within each hand of the biodiversity clock enhances its dimensionality and information content.

The proportion of endemic and keystone species within each of the three conservation classes at KSA and EDK are represented in the corresponding segment of hour, minute, and seconds hand, respectively. *Artocarpus hirsutus, Holigarna arnottiana, Hydnocarpus pentandra,* and *Hopea ponga, Syzygium zeylanicum, Cinnamomum malabatrum* are few of the endemic tree species observed in KSA and EDK, respectively. Sacred groves are known refuge of endemic species [30, 39]. The biodiversity clock of KSA and EDK revealed a higher proportion of endemic trees in vulnerable and endangered classes. More than half of the vulnerable and endangered tree species in KSA and EDK were endemics. Endemic species convey information about the uniqueness of habitat [40, 41], environmental quality [42], and biogeography [41]. The prominent depiction of the presence of endemic species in vulnerable (VU) and endangered (EN) classes in the biodiversity clock, can be used as weights while prioritizing conservation planning to arrest local extinction [43, 44]. The weight (relative importance) of a site will increase if its biodiversity clock bears a + symbol in the seconds-hand (EN), as compared to that of a site with + symbol on the hour-hand (LC).

The presence of keystone species paints a picture of the structure, composition, and function of the ecosystem. Keystones are exceptional species, which help to shape the community structure and maintain diversity [45]. The disproportionately large influence exerted by keystone species [46], enhances their conservation importance [47]. Removing or adding keystone species to a community will trigger a cascade of interactions that can have drastic effects on the ecosystem [48]. *Ficus religiosa* and *Ficus benghalensis* were the keystone tree-species observed in KSA and EDK, respectively. Both *Ficus religiosa* and *Ficus benghalensis* are LC tree species. The Genus *Ficus* is an important food source for ecologically important frugivores [49, 50]. *Ficus benghalensis*. in sacred groves plays the role of a keystone species providing a niche for a large number of birds and plants [51]. The occurrence of keystone species in VU or EN class can be taken as a surrogate of ecosystem fragility.

Biological invasion is a creeping disaster [52]. Invasive species are quick to cash-in on the degradation of habitat, which is often associated with anthropogenic interference [53, 54]. Their presence is a reflection of the quality of habitat. Human-induced fragmentation of landscape results in canopy gaps that alters the light regime, and act as gateways of plant invasion [55]. The solid circle on the dial of the biodiversity clocks in both the sites indicate dense and evenly distributed canopy. Closed canopy at both the sacred groves kept invasive species beyond their respective boundaries. However, *Mikania micrantha*, one of the high-risk invasive plant species reported from Kerala, India [56], was found lurking in the fringe of KSA. We attribute this to the human activities in the adjoining plantation area. Ground-based measurement of the canopy gap (*CNG*) complements the remote estimation of forest cover (Aichi target 5). It assumes importance in fine-resolution biodiversity monitoring that is beyond spatial resolvability and visibility constraints of space-based sensors.

### Conservation Triangle

The three dimensions of the IUCN conservation classes can be denoted without loss of information in the conservation triangle. It complements the biodiversity clock. Every point within the conservation triangle corresponds to a unique combination of the proportion of the three IUCN conservation classes (LC,VU, and EN). The position of a site within the conservation triangle is a surrogate that can be leveraged for effective prioritization of conservation intervention. The conservation locus of a site with an equal proportion of the three IUCN conservation classes will fall at the centroid of the triangle. With an increasing proportion of endangered species (*PEN*), the loci migrate upwards towards the apex. Similarily the conservation locus will shift to either the lower right or lower left vertices, corresponding to the increase in respective proportion of least concern (*PLC*) and vulnerable (*PVU*) species in the population. The relatively higher proportion of endangered (*PEN*) and vulnerable (*PVU*) tree species at EDK shifted its conservation locus away from the LC edge of the conservation triangle, as compared to that of KSA.

### Biodiversity Clock and Conservation Triangle: Tools for Conservation Target

In the concluding months of the international decade of dedicated, legally binding action on the global biodiversity framework, we are staring at a half-filled cup. Albeit an enhanced awareness of the importance of biodiversity [22, 57], the goals remain afar [58]. More *Parties* have enacted legislation and are striving to meet their national commitments. However, at a global level, the efforts at monitoring biodiversity changes continue to remain fragmented and are too modest. Our knowledge about biodiversity mostly stems from discrete research projects that have non-overlapping themes [22, 59]. While across the conservation spectrum, there is a strong focus to protect habitats in order to conserve biodiversity, we know little about their effectiveness [60]. With slightly less than 15% global terrestrial habitats protected [61], and even lesser area under continuous *proximate* monitoring; our prophecies on trends in biodiversity rests on a weak foundation. Pooling the existing biodiversity monitoring programs and initiating intelligent BIodiversity Observation Sites (BIOS) to ensure representative global sampling will reinforce the foundation and pave the way for the post-2020 global biodiversity framework.

The biodiversity clock and conservation triangle introduced in this paper posit as ideal tools that act as effective convergence plane to represent biological diversity and streamline conservation efforts. While the former denotes the compositional, structural, and functional aspects of biodiversity, the latter is invaluable to prioritize conservation investment. Biodiversity, represented as time on a dial, is a powerful means to convey information to the general public. Researchers can read the angular values of the clock hands to arrive at precise information for effective policymaking. Shannon index of the diversity of the site, the abundance of invasive species, presence of endemic, and the number keystone species, the % canopy cover, and the physiography are the other dimensions that can be directly read from the biodiversity clock. Diving deeper, researchers can use the information conveyed in the clock and the database to estimate multiple indices, *viz.* Simpson Index, Simpson Effective number, Simpson Evenness [62], Brillouin index [63], Margalef Index [64], Pielou’s index [65], Berger-Parker index [66], Hill Number [67], Menhinick’s index [68], Rao’s quadratic entropy [69] and Clarke and Warwick’s taxonomic distinctness index [70, 71].

The scale independence of the biodiversity clock makes an ideal gateway to harness a huge volume of biodiversity data generated within academia that currently remain outside the formal framework [72, 73]. In addition, the biodiversity clock and conservation triangle concurrently fill the gaps in indicators of multiple Aichi targets and SDGs. The universality of clock reading renders the unparalleled advantage to make the biodiversity clock an effective representation to increase public awareness of biodiversity (target 1). In this case, the biodiversity clock explicitly represents 18 dimensions in a two-dimensional frame. The lucid pictorial representation of biodiversity as *time* exemplifies persuasive communication, which is a proven strategy to frame messages that trigger behavioral changes [74]. Together the clock and the conservation triangle are one step ahead of the biodiversity dashboard [57].

The ecophysical dimension of canopy openness (*CNG*) represented in the biodiversity clock can complement as an indicator of tree cover, especially at finer scales and in isolated patches of vegetation (target 5). It can further be effectively used to downscale tree cover trends in finer resolution. The dimension of invasive species embedded in the biodiversity clock can act as a framework to understand trends in the distribution and population of invasive species. (target 9). Since canopy gaps are known gateways of plant invasion, read together with the invasive species, it can be a surrogate for studying the prospective trend of invasion. Over time, the biodiversity clock will constitute a robust inventory of biodiversity (target 19). In addition, the clock with its constituents gives a good ecological representation of a site (target 11). The conservation triangle directly feeds into Aichi targets 1 and 11. The conservation locus paints a picture of the state of a site, and its temporal changes form an objective metric to assess the effectiveness of the management of conserved areas.

The biodiversity clock and conservation triangle seek to harness the potentials of emerging technologies towards the 2050 vision for biodiversity. We envisage it as an open framework that acts as a convergence plane for the ongoing independent efforts. The clock as an indicator of biodiversity is a novel means that conveys holistic information on biodiversity. A cloud-mounted biodiversity clock can assimilate data sans boundaries and act as a framework to monitor the status, trends, and share data. It can act as guard rails, which will help us to be on track to the biodiversity targets [22]. Integrated with the global programs and with pre-selected *Specific, Measurable, Achievable, Realistic*, and *Time-specific* (SMART) objectives [75], the system can reveal hitherto unseen trends in biodiversity and even advise preemptive interventions.

## Conclusion

Inherent multi-dimensionality of biodiversity makes it difficult to represent it in a form that is uniformly cognized. The biodiversity clock ingeniously leverages the universal homogeneity of clock reading to express plant diversity. The utility of the clock representation was exemplified as a case study in two sacred groves in Kerala, India. Loci of each grove within the conservation triangle are indicative of its conservation status. The clock and triangle posit as an effective tool to address multiple biodiversity targets. The framework has the potential to leverage the prospects of emerging technologies and act as a travelator to post the 2020 global biodiversity framework.

## Acknowledgments

We thank Dr. Saji Gopinath, Director, Indian Institute of Information Technology and Management – Kerala for providing all necessary support to carry out the study. Thanks are also due to the custodians of the two sacred groves, Kavil Shree Maheswar Ashramam, and Edayilekkadu Kavu, Kerala, India, to facilitate the survey.

## Supporting information

**S1 File. Spreadsheet template for ground data collection**

**S2 File. Description of components of the biodiversity clock**

**S3 File. R code for Conservation Triangle**

**S4 File. List of tree species, in the two sacred groves EDK and KSA along with their conservation status**

